# Tethered NOS-3, a nematode Nanos RNA-binding protein, enhances reporter expression and mRNA stability

**DOI:** 10.1101/2021.01.24.428005

**Authors:** Jonathan Doenier, Tina R Lynch, Judith Kimble, Scott T Aoki

## Abstract

Robust methods are critical for testing the *in vivo* regulatory mechanism of RNA binding proteins. Here we report improvement of a protein-mRNA tethering assay to probe the function of an RNA binding protein in its natural context within the *C. elegans* adult germline. The assay relies on a dual reporter expressing two mRNAs from a single promoter and resolved by trans-splicing. The *gfp* reporter 3’UTR harbors functional binding elements for λN22 peptide, while the *mCherry* reporter 3’UTR carries mutated nonfunctional elements. This strategy enables internally controlled quantitation of reporter protein by immunofluorescence and mRNA by smFISH. To test the new system, we analyzed a *C. elegans* Nanos protein, NOS-3, which serves as a post-transcriptional regulator of germ cell fate. Unexpectedly, tethered NOS-3 enhanced reporter expression. We confirmed this enhancement activity with a second reporter engineered at an endogenous germline gene. NOS-3 enhancement of reporter expression was associated with its N-terminal intrinsically disordered region, not its C-terminal zinc fingers. RNA quantitation revealed that tethered NOS-3 enhances stability of the reporter mRNA. We suggest that this direct NOS-3 enhancement activity may explain a paradox: classically Nanos proteins are expected to repress RNA, but *nos-3* had been found to promote *gld-1* expression, an effect that could be direct. Regardless, the new dual reporter dramatically improves *in situ* quantitation of reporter expression after RBP tethering to determine its molecular mechanism in a multicellular tissue.

## Introduction

Protein-mRNA tethering is a well-established method to investigate the direct regulatory effects of RNA binding proteins (RBPs). These assays rely on two components: an RBP tagged with a λN22 peptide or MS2 coat protein domain and a reporter mRNA harboring binding sites for λN22 or MS2 in its 3’UTR (Baron-Benhamou et al. 2004; Coller and Wickens 2007). The RBP is thus tethered to reporter mRNA with high affinity and specificity, and its regulatory effect inferred from changes in expression or stability of the reporter RNA compared to a control. Tethering assays have proven tremendously useful. They have revealed that some RBPs downregulate mRNA expression by promoting RNA turnover (e.g. (Bhandari et al. 2014; Raisch et al. 2016)) or repressing translation (e.g. (Pillai et al. 2004), while others upregulate mRNA expression by stabilizing RNA (e.g. (Coller et al. 1998; Gray et al. 2000)) or enhancing translation (e.g. (De Gregorio et al. 1999)). Tethering assays are thus an exceptional tool to dissect the molecular functions of RBPs.

The regulatory effect of a tethered RBP is deduced from measurements of both the reporter RNA’s stability and translation. When assays are done in cultured cells or yeast, bulk detection methods are sufficient, but when conducted in a multicellular tissue or organism, cell-specific methods become essential. Here we report the development of a new tethering assay for use in the *C. elegans* germline, a richly patterned tissue in which mRNAs are dynamically regulated as cells develop from dividing stem cells to differentiating gametes (Hubbard and Schedl 2019). This assay takes advantage of a dual reporter that includes an internal control for better quantitation, a PEST domain to restrict reporter protein half-life, and high affinity epitope tags for each reporter protein. We test our assay with NOS-3, an RBP belonging to the conserved Nanos family (Kraemer et al. 1999; Subramaniam and Seydoux 1999). Nanos RBPs regulate germline development from nematodes to humans (Tsuda et al. 2003; Suzuki et al. 2012; Kusz-Zamelczyk et al. 2013). Tethering assays of human and *Drosophila* Nanos orthologs, performed in HEK293 or S2 cells respectively, demonstrate that both are repressors of reporter expression and identified a Not1-interacting domain responsible for that repression (Bhandari et al. 2014; Raisch et al. 2016). We selected NOS-3 to test our new assay, and expected it would repress reporter expression. However, we found instead that tethered nematode NOS-3 enhances reporter expression, a surprising result that implies that NOS-3 may promote expression of its target mRNAs in the *C. elegans* germline.

## Results and Discussion

We set out to improve the tethering assay in the *C. elegans* germline. Our starting point was a λN22-boxB tethering system previously used in this tissue (Wedeles et al. 2013; Aoki et al. 2018; Aoki et al. 2019). The earlier system tagged the RBP of interest with λN22 peptide, henceforth λN, and recruited RBP∷λN to germline-expressed, reporter mRNA via boxB RNA hairpins in its 3’ UTR (Aoki et al. 2018; Aoki et al. 2019) (**Figure 1A**). λN binds to boxB hairpins with high affinity and specificity (Baron-Benhamou et al. 2004). The GFP-tagged, histone H2B reporter protein localizes to the nucleus of the expressing cell for straightforward visualization. While functional, the previous system lacked internal controls to compare expression between animals. Histone reporters can also persist and mark cells or their daughters, expanding perceived protein expression boundaries (Merritt et al. 2008). Thus, more accurate measurement of reporter expression required additional revisions to the assay.

**Figure 1.**
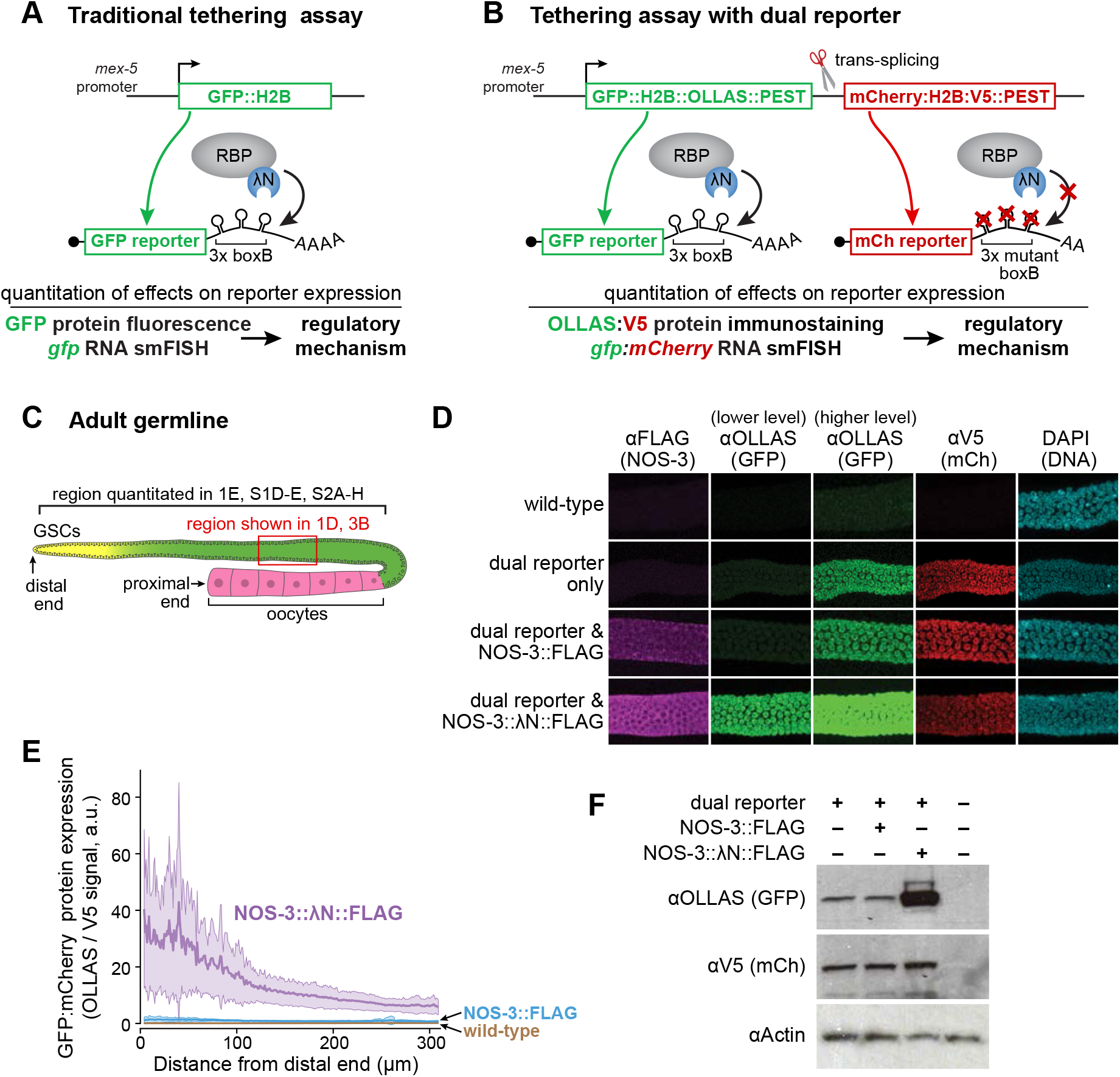
Tethered NOS-3 increases GFP reporter expression. **(A)** Previous tethering assay used in the *C. elegans* germline (Aoki et al. 2018; Aoki et al. 2019). GFP reporter mRNA is under control of a germline-expressed, *mex-5* promoter and has three boxB stem loops in its 3’UTR. The RNA-binding protein (RBP) is tagged with λN. **(B)** Dual reporter tethering assay (this work). The nascent transcript driven by *mex-5* promoter is resolved by trans-splicing into two mRNAs that encode distinct reporters. The *gfp* reporter RNA has three functional boxB stem loops in its 3’UTR; the *mCherry* reporter 3’UTR has three mutated boxB stem loops that do not bind λN and therefore provides an internal control. **(C)** Schematic of *C. elegans* germline. Germline stem cells (GSCs) reside in the progenitor zone (yellow); GSC daughters enter and progress through meiotic prophase (green) and finally differentiate into oocytes (pink); red box, mid-pachytene region imaged in **Figure 1D**; bracket, region quantitated in several figures begins at distal end and goes 310 μm, stopping near proximal end of the pachytene region. **(D)** Confocal images (max projection) of germlines stained to detect NOS-3∷FLAG and NOS-3∷λN∷FLAG (λFLAG, column 1); GFP∷H2B∷OLLAS∷PEST (λOLLAS, columns 2 and 3); V5 antibodies to detect mCherry∷H2B∷V5∷PEST (λV5, column 4) and DNA (DAPI, column 5). Images in each column were visualized at the same level in Image J; the column 2 OLLAS level was optimized for reporter expression with NOS-3∷N∷FLAG, while the column 3 level was optimized for expression with the dual reporter only and NOS-3∷FLAG. Wild-type has no tagged proteins and serves as a negative control. Note that images for wild-type, reporter only, NOS-3∷FLAG and NOS-3∷λN∷FLAG are replicated in **Figure 3B**. **(E)** In situ quantitation of GFP (λOLLAS) normalized to mCherry (λV5) reporter proteins, as a function of germline position. Lines show averages and shading shows one standard deviation (see Methods). Wild-type serves as a negative control. Wild-type (n=54), NOS-3∷FLAG (n=34), NOS-3∷λN∷FLAG (n=52) worms analyzed. Same curves and values are included in **Figure 3D, Figure S1D-E** and **Figure S2**. **(F)** Immunoblot to assay GFP (λOLLAS) and mCherry (λV5) reporter protein expression. Actin serves as a loading control, and wild type worms as the negative control (last lane).

The improved reporter incorporates three new components into the previous system, all with the goal of enhancing quantitation of expression in specific cell types (**Figure 1B** and **Figure S1A)**. First, we generated a reporter operon that introduces a second mRNA reporter as an internal control, as conceived by others (Merritt et al. 2008). A single nascent transcript generates two reporter mRNAs by trans-splicing: one encodes GFP-H2B and can bind RBP∷λN via boxB RNA hairpins in its 3’ UTR; the second encodes mCherry-H2B and cannot bind RBP∷λN because mutated boxB RNA hairpins in its 3’ UTR abrogate λN binding (Chattopadhyay et al. 1995). Second, we added a PEST domain to the C-terminus of both reporter proteins to shorten half-life of the fusion protein and restrict it to cells expressing reporter RNA (Frand et al. 2005; Farley and Ryder 2012; Kaymak et al. 2016). Third, we inserted high affinity epitope tags with commercially available antibodies: OLLAS into GFP∷H2B∷PEST, and V5 into mCherry∷H2B∷PEST. A construct encoding the dual reporter was integrated into the *C. elegans* genome as a Mos single copy insertion (MosSCI, see Methods). The *C. elegans* germline is patterned along its U-shaped distal-proximal axis with germline stem cells (GSCs) at the distal end, progressive differentiation of their daughters more proximally, and differentiated gametes at the proximal end (Hubbard and Schedl 2019) (**Figure 1C**). Immunostaining against OLLAS and V5 tags revealed that both GFP and mCherry histone H2B reporter proteins expressed and co-localized with DNA in germ cell nuclei (**Figure 1D**). This dual reporter thus allows for precise quantitation of expression from both tethered and untethered mRNA in individual cells within the *C. elegans* germline (**Figure 1B**).

We tested the dual reporter with NOS-3, an mRNA binding protein and germ cell fate regulator. NOS-3 regulates two fate choices that occur along the germline axis, the sperm/oocyte and mitosis/meiosis decisions (Kraemer et al. 1999; Eckmann et al. 2004; Hansen et al. 2004). However, genetic analyses of how NOS-3 functions in those two decisions poses a paradox. NOS-3 decreases expression of FEM-3 (Arur et al. 2011), a positive regulator of the sperm fate, but increases expression of GLD-1 (Hansen et al. 2004; Brenner and Schedl 2016), a positive regulator of meiotic entry. However, these genetic results do not address whether NOS-3 regulation is direct or indirect. If NOS-3 were primarily a repressor, like other members of the Nanos family (see Introduction), its enhancement of GLD-1 expression might be indirect, perhaps by repressing a *gld-1* mRNA repressor.

To ask how NOS-3 regulates expression when tethered, we inserted a 3xFLAG epitope tag with or without λN into the endogenous *nos-3* gene, using CRISPR/Cas9 gene editing (**Figure S1B**). Both NOS-3∷FLAG and NOS-3∷λN∷FLAG proteins were expressed in the germline cytoplasm (**Figure 1D, first column**), as reported previously for endogenous NOS-3 (Kraemer et al. 1999). Moreover, the λN∷FLAG-tagged NOS-3 behaved like wild-type NOS-3 in a genetic assay (**Figure S1C)**, validating its use to test NOS-3 function. We proceeded to ask how tethered NOS-3 affects reporter expression. Expression of GFP and mCherry reporters was assayed by immunostaining against their OLLAS and V5 tags, respectively. When OLLAS staining was optimized for GFP reporter expression in the strain carrying NOS-3∷λN∷FLAG, the expression was only faintly detectable in germlines with the dual reporter only or the reporter plus NOS-3∷FLAG (**Figure 1D, second column**). However, when optimized for expression in the strains carrying the dual reporter only or the reporter plus NOS-3∷FLAG, expression was easily observed in those germlines but saturated with NOS-3∷λN∷FLAG (**Figure 1D, third column**). Thus, addition of the λN tag to NOS-3 dramatically increased GFP reporter expression (**Figure 1D, second and third columns,** and **Figure S1D**). By contrast, expression of the mCherry internal control reporter protein was observed at comparable levels in all germlines carrying the dual reporter (**Figure 1D, fourth column,** and **Figure S1E**). We used this mCherry control to normalize for variable reporter protein expression across worms (**Figure 1B**). Ratios of GFP to mCherry reporter expression were quantified as a function of germline position (**Figure 1E**; see **Figures S1D-E** for graphs of the two reporter proteins on their own). In NOS-3∷FLAG germlines, the ratio was modest, but in NOS-3∷λN∷FLAG germlines, the ratio increased significantly throughout the distal germline arm. Immunoblots of the reporter proteins confirmed enhancement of GFP reporter expression specifically (**Figure 1F).** We conclude that NOS-3 can enhance reporter expression when tethered.

The NOS-3 enhancement of reporter expression was unexpected because other Nanos homologs repress expression when tethered (Bhandari et al. 2014; Raisch et al. 2016). To further test the effect of tethered NOS-3, we engineered an endogenous germline gene to function as a tethering reporter. Three boxB hairpins were inserted into the 3’UTR region of the *pgl-1* gene, which encodes a SNAP-tagged PGL-1 (**Figure 2A**). The SNAP tag permits straightforward visualization of PGL-1 protein in P granules without affecting its function (Aoki et al. 2019). As with the dual reporter, NOS-3∷λN∷FLAG dramatically increased PGL-1∷SNAP abundance when compared to NOS-3∷FLAG (**Figure 2B, C**). We conclude that tethered NOS-3 enhances expression of two distinct reporter mRNAs.

**Figure 2.**
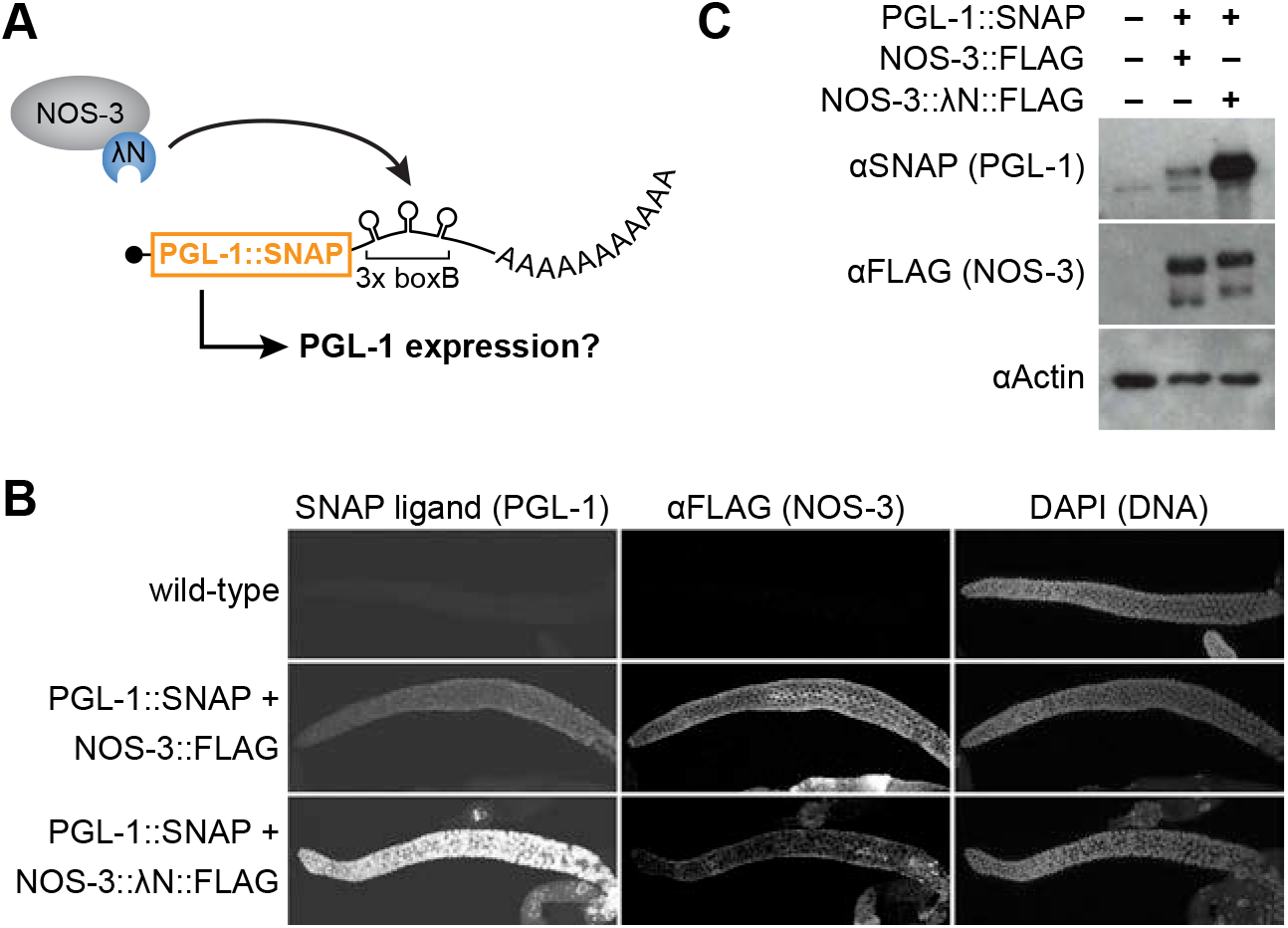
Tethered NOS-3 increases expression of *pgl-1* endogenous gene. **(A)** Tethering assay using SNAP-tagged *pgl-1* gene as the reporter. Endogenous *pgl-1* includes a SNAP tag to visualize its protein product and 3x boxB hairpins to recruit NOS-3∷N∷FLAG to its 3’UTR. **(B)** Tethered NOS-3 increases expression of PGL-1 protein in the worm germline. Confocal images (max projection) of adult gonads stained for PGL-1∷SNAP (SNAP ligand), NOS-3 (⍺FLAG), and DNA (DAPI). Wild-type worms serve as a negative control. **(C)** Immunoblot to assay PGL-1 (⍺SNAP) and NOS-3 (⍺FLAG) protein expression. Actin serves as the loading control, and wild-type worms as a negative control (first lane).

To probe the region or regions within NOS-3 responsible for enhancing expression, we generated a battery of NOS-3∷λN∷FLAG variants by CRISPR/Cas9 gene editing of the endogenous locus. Full length NOS-3 protein consists of a large N-terminal region predicted to be intrinsically disordered (analyzed by IUPred2A (Erdos and Dosztanyi 2020)), and a Nanos-like zinc finger near the C-terminus (Kraemer et al. 1999; Subramaniam and Seydoux 1999) (**Figure 3A, top**). Variants were made with in-frame deletions or a premature stop codon (**Figure 3A, bottom**). All protein variants described were expressed, albeit at differing expression levels (**Figure 3B, first column; 3C, bottom panel**). Reporter expression was examined in all variants by imaging (**Figure 3B**) and calculating ratios of GFP to mCherry reporter protein abundance in germlines (**Figure 3D, Figure S2**), as well as by immunoblot (**Figure 3C**). All variants enhanced GFP reporter expression relative to the internal control to some extent (**Figure 3D, Table S1, Figure S2**). Of note, the enhancement does not require the C-terminal Zinc finger but instead relies on the intrinsically disordered region constituting the large N-terminus (**Figure 3D)**. Intriguingly, this region also includes the FBF-1 interaction domain (Kraemer et al. 1999) and ERK/MAP Kinase (MPK) docking site (Arur et al. 2011). Regardless, our results suggest that enhancement activity may be distributed across NOS-3 protein.

**Figure 3.**
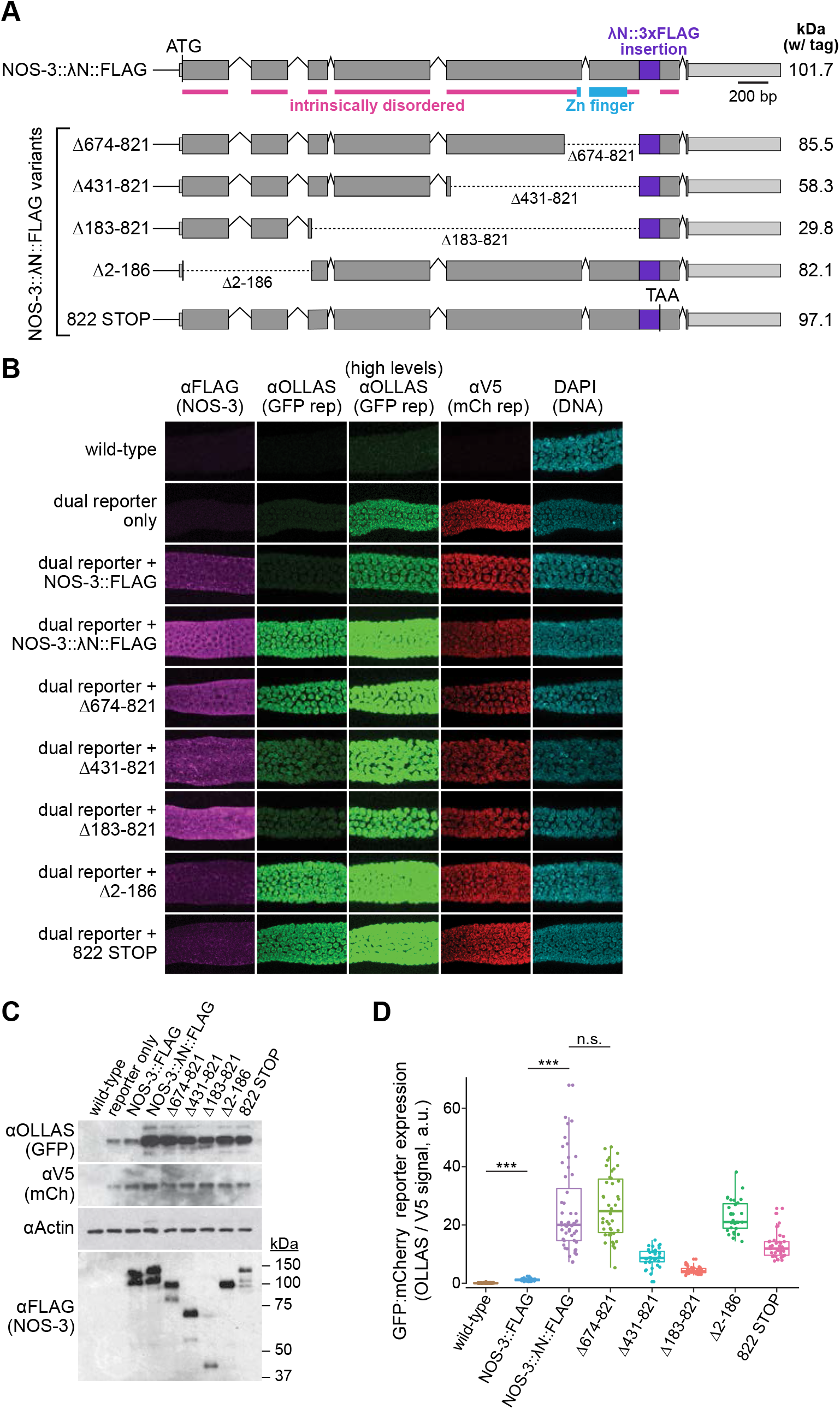
Analysis of tethered NOS-3 deletion mutants for enhancement of reporter expression. **(A)***nos-3* locus and variants used to map enhancement activity. Variants include four in-frame deletions and a premature STOP codon that removes amino acids C-terminal to the tag insert. Variant protein sizes (kilodaltons, kDa) shown at right. Coding regions, grey; untranslated regions, light grey; λN∷FLAG insert, purple. Intrinsically disordered regions were predicted by IUPred2A (Erdos and Dosztanyi 2020). **(B)** Confocal images (max projection) of adult germlines. Conventions as detailed in **Figure 1D**. See **Figure 1B** for germline image location. Note that images for wild-type, reporter only, NOS-3∷FLAG and NOS-3∷λN∷FLAG are replicated from **Figure 1D**. **(C)** Immunoblot to assay expression of reporter proteins (above) and NOS-3 variant proteins (below). Conventions as in **Figure 1F**. SDS-PAGE sizes (kDa) shown to right for the NOS-3 immunoblot. **(D)** Relative abundance of GFP and mCherry reporter proteins. Ratio of signals from GFP (⍺OLLAS) and mCherry (⍺V5) reporter proteins, averaged across distal 100 μm of germline and normalized to wild-type negative control. Wild-type (n=54), NOS-3∷FLAG (n=34), NOS-3∷λN∷FLAG (n=52), λ183-821 (n=37), λ431-821 (n=36), λ674-821 (n=42), λ2-186 (n=31), 822 STOP (n=37) worms analyzed. Curves and values of wild-type, NOS-3∷FLAG and NOS-3∷N∷FLAG are same as reported in **Figure 1E** and **Figure S2**. ***, p-value <0.0001. n.s., not significant (p-value=0.5735). All p-values for comparisons between strains can be found in **Table S1**.

To investigate how tethered NOS-3 enhances expression of the *gfp* reporter RNA, we assessed the abundance of the two dual reporter mRNAs in germlines harboring NOS-3∷λN∷FLAG or controls. The *gfp* and *mCherry* reporter mRNAs were visualized using distinct smFISH probe sets, which permitted simultaneous imaging of both reporter mRNAs in the same animal (**Figure 4A;** see Methods). We predicted that tethered NOS-3 affects either stability or translation of the *gfp* reporter mRNA. If tethered NOS-3 stabilizes its target RNA, the *gfp* mRNA levels should be higher in animals carrying NOS-3 tagged with λN than those without (**Figure 4B, top**). If tethered NOS-3 solely promotes translation, the *gfp* mRNA levels should be similar with or without λN (**Figure 4B, bottom**). *mCherry* mRNA levels serve as a normalizing control for transcription.

**Figure 4.**
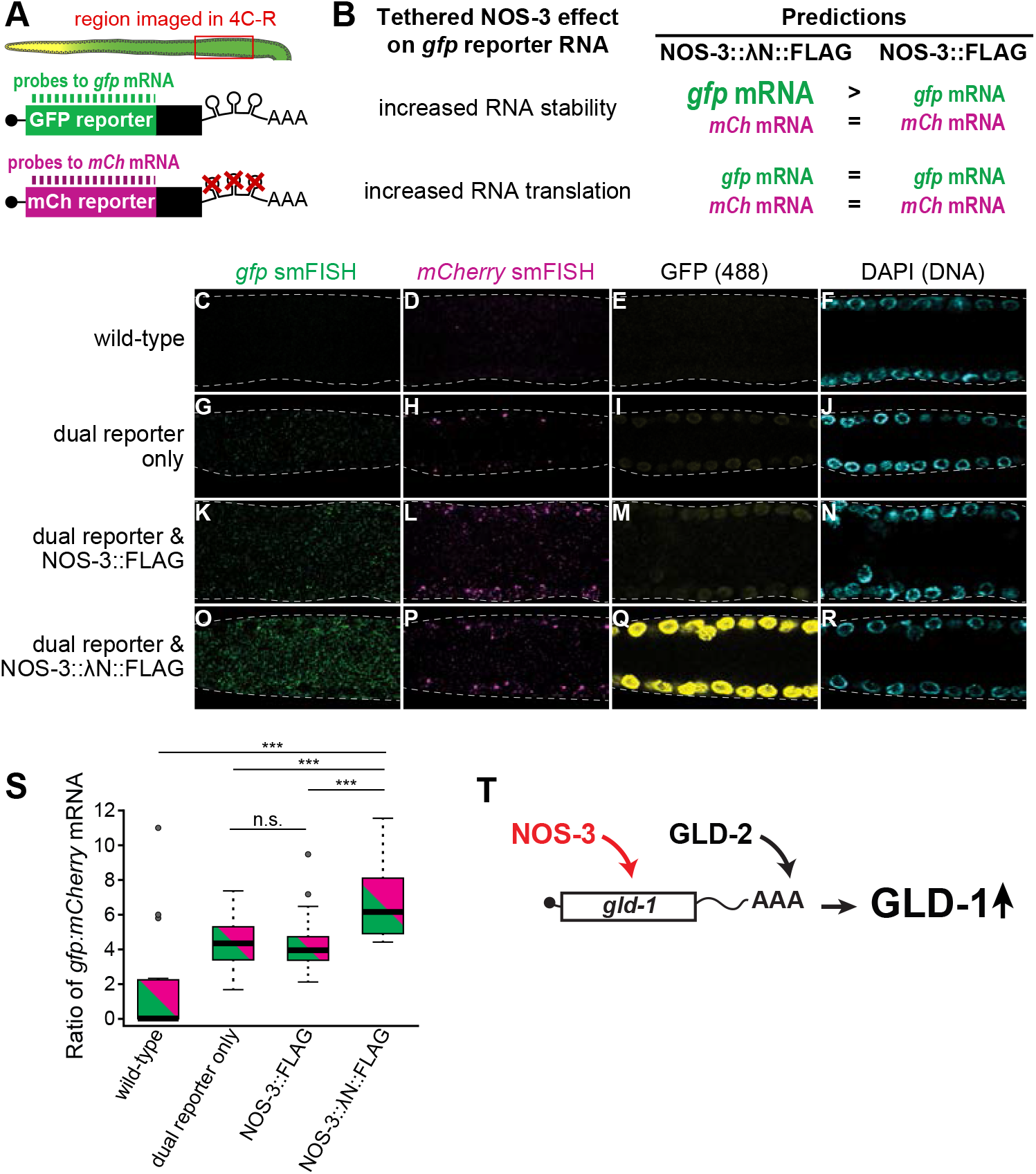
Tethered NOS-3 enhances stability of *gfp* reporter mRNAs. **(A)** Experimental design. Reporter mRNAs from dual reporter visualized in dissected gonads (top) with differentially labeled smFISH probes to *gfp* and *mCherry (mCh)* mRNAs (bottom). **(B)** Tethered NOS-3 might increase reporter expression by affecting either stability or translation of the reporter RNA. These two mechanisms have distinct predictions for effects on reporter mRNA abundance. More or less RNA abundance is depicted in the figure as larger or smaller letters, respectively, in animals possessing tethered NOS-3∷λN∷FLAG (left column) or untethered NOS-3∷FLAG (right column). **(C-R)** Representative images of smFISH in wild-type, reporter only, NOS-3∷FLAG and NOS-3∷λN∷FLAG germlines. Images are average projections of 5 slices (0.3 μm) from confocal microscopy. *gfp* mRNAs (*gfp* smFISH, green), *mCherry* mRNAs (*mCherry* smFISH, magenta), GFP protein (fluorescence signal (488), yellow) or DNA (DAPI, light blue). **(S)** Ratios of *gfp:mCherry* mRNAs from smFISH quantitation. ***, p-value<0.0001. n.s., not significant (p-value=0.6352). All reporter strains were significantly different than wild-type (p-value<0.0001). **(T)** Model that NOS-3 directly enhances expression of *gld-1* mRNA, perhaps together with the GLD-2 poly(A) polymerase.

smFISH detected *gfp* and *mCherry* mRNAs in all germlines carrying the dual reporter (**Fig 4G,H; 4K,L; 4O,P),** but not in wild-type germlines without the reporter (**Fig 4C-D**). mRNA signals appeared as cytoplasmic spots, as established in other studies (Raj et al. 2008; Lee et al. 2017). The *gfp* mRNAs were more abundant than *mCherry* mRNAs (**Figure 4G,H**), which could result from differences in probe binding efficiency or differential mRNA processing. Fluorescence of the GFP protein was higher in reporter expressing germlines with NOS-3 tagged with λN (**Figure 4M** versus **Figure 4Q**), consistent with immunostaining of its epitope tag. The number of *gfp* mRNAs was normalized to *mCherry* mRNA counts in the same germlines, and *gfp:mCherry* mRNA abundance ratios were compared between strains (**Figure 4S**). This analysis revealed that the normalized *gfp* mRNA levels were significantly different between tethered and untethered NOS-3 (**Figure 4S**). As expected, all germlines carrying the dual reporter had more detectable reporter mRNA signal than the wild-type control. More importantly, the *gfp:mCherry* mRNA ratios were similar for NOS-3∷FLAG and wild-type NOS-3, but increased for tethered NOS-3∷λN∷FLAG (**Figure 4S**). This result fits the prediction that NOS-3 enhances *gfp* expression by increasing mRNA stability, though we cannot exclude an additional effect on enhancing translation.

The enhancement activity of tethered NOS-3 was surprising given that its homologs act as repressors when tethered (Bhandari et al. 2014; Raisch et al. 2016). Yet the NOS-3 enhancement activity may help explain a puzzling genetic result. Removal of NOS-3 increases FEM-3 protein abundance (Arur et al. 2011), consistent with repressive activity, but its removal also lowers GLD-1 protein abundance (Hansen et al. 2004; Brenner and Schedl 2016). One explanation for the GLD-1 decrease, based on dogma that Nanos proteins are repressors, is that NOS-3 represses a *gld-1* repressor and the GLD-1 change is indirect. However, here we find that tethered NOS-3 directly enhances expression of two distinct mRNAs, the *gfp* mRNA of the dual reporter and the endogenous *pgl-1* mRNA that we engineered for tethering. We therefore suggest that NOS-3 likely promotes GLD-1 expression directly (**Figure 4T**). NOS-3 enhancement may work in parallel with the GLD-2 poly(A) polymerase, which also increases expression of *gld-1* mRNA through elongation of its polyA tail (Suh et al. 2006; Suh et al. 2009). We emphasize that a NOS-3 repressive activity remains possible. MPK phosphorylation may have modulated a switch between NOS-3 acting as a repressor and activator (Arur et al. 2011). However, that MPK phosphorylation is spatially restricted to one region of the germline, whereas NOS-3 enhancement activity is not similarly restricted. Clearly, further investigation of NOS-3 enhancement activity is warranted as is investigation of classical Nanos orthologs in their natural context.

The enhancement activity of tethered NOS-3 also expands the genetic toolkit for analyzing protein function. Methods to decrease protein abundance are readily available with classical genetics, RNAi and gene editing, but methods to increase protein abundance have been less robust, though examples exist. Expression of specific genes can be increased by modifying CRISPR/Cas9 (Konermann et al. 2015) or tethering proteins associated with translation, like polyA binding protein (PABP) (Coller et al. 1998), eIF4E (De Gregorio et al. 2001) or eIF4G (De Gregorio et al. 1999). We can now add NOS-3 tethering as a method to increase gene expression, at least in the *C. elegans* germline. More broadly, pairing the dual reporter tethering assay with smFISH allows direct investigation of mechanistic functions of RNA binding proteins in complex multicellular tissues. By analyzing the ratios of tethered versus untethered transcripts, one can assess mechanisms of mRNA turnover and translational modulation. This mechanistic dissection performed within the natural context of a multicellular organism allows regulatory mechanisms to be analyzed in different tissues and cells and at different stages of development. The dual reporter tethering-smFISH pairing is adaptable to other organisms with single gene editing and shows promise to elucidate the post-transcriptional regulatory mechanisms of a wide variety of mRNA binding proteins.

## Abbreviations

RNA: ribonucleic acid;
UTR: untranslated region
GFP: green fluorescent protein
CRISPR: Clustered Regularly Interspaced Short Palindromic Repeats
RNAi: ribonucleic acid interference
smFISH: single molecule Fluorescence In Situ Hybridization

## Acknowledgements

We thank L. Lavis for supplying the SNAP JF 549 ligand, C. Lee for smFISH advice, L. Vanderploeg for help with figure revisions, and members of the Kimble and Wickens Labs for helpful discussions. JD and TRL conceived and performed the experiments and helped write the paper; JK and STA conceived the experiments and wrote the paper. TRL was supported by the NSF; this material is based upon work supported by the National Science Foundation Graduate Research Fellowship under Grant No. DGE-1747503. Any opinions, findings, and conclusions or recommendations expressed in this material are those of the authors and do not reflect the views of the National Science Foundation. JK was an Investigator of the Howard Hughes Medical Institute and is now supported by NIH R01 GM134119. STA was supported by NIH K99 HD081208.

## Methods

### Worm maintenance and strains

Strains were maintained at 20° or 25°C, as described (Brenner 1974). **Table S2** lists strains analyzed, which are available upon request.

### CRISPR-Cas9 Gene Editing

CRISPR-Cas9 genome editing was used to modify endogenous *nos-3* and *pgl-1* gene loci, following an established protocol (Paix et al. 2015). Briefly, worms were injected with a ribonucleoprotein complex consisting of recombinant Cas-9 protein, tracrRNA, gene-specific crRNAs, and a single stranded DNA repair template. The injection mix included a crRNA and repair template designed to induce a dominant loss of function mutation in *unc-58* as a marker for successful editing (Arribere et al. 2014). Unc progeny from injected animals were singled and PCR screened for edits at the intended locus. Animals homozygous for edits were isolated and proper editing confirmed by Sanger sequencing; confirmed homozygotes were outcrossed with N2 a minimum of two times before analysis. **Table S3** lists guide RNAs and repair DNA oligos.

### MosSCI insertion of the dual reporter gene

The dual reporter construct included a *mex-5* promoter (Merritt et al. 2008), a region encoding eGFP∷Histone H2B tagged with a mouse ornithine decarboxylase (MODC) PEST domain (Li et al. 1998) and a 1xOLLAS epitope tag (Park et al. 2008), a *tbb-2* 3’UTR harboring a 3x boxB (Wang et al. 2011), a *gpd-2/gpd-3* transplice site (Huang et al. 2001), an mCherry∷Histone H2B fusion tagged with MODC PEST and 1xV5 epitope tag (Hanke et al. 1992), a *tbb-2* 3’UTR harboring a mutated 3x boxB sequence, and a *tbb-1* intergenic region (**Figure S1A**). This construct was inserted into the *C. elegans* genome as a Mos1-mediated single copy transgene insertion (MosSCI) (Nance and Frokjaer-Jensen 2019). To this end, plasmids encoding the Mos1 transposon, injection markers, and dual reporter were injected into the gonad of EG8081 worms (Frokjaer-Jensen et al. 2008) containing the targeted oxTi177 site on chromosome IV. Progeny of injected animals were screened for integration of the dual reporter and loss of extrachromosomal arrays with injection marker. The inserted construct was verified by Sanger sequencing. Animals with the insertion were outcrossed two times and propagated as homozygotes at 25°C, to enhance reporter expression.

### Gonad immunostaining, imaging and fluorescence quantitation

Germline staining was performed following standard protocols (Crittenden et al. 2017). For immunostaining, gonads were extruded and fixed. In **Figures 1, 3, and 4**, and **Figure S1 and S2**, gonads were fixed with 1% paraformaldehyde in 100 mM K_2_HPO_4_ for 20 minutes and permeabilized with ice-cold methanol for 10 minutes. For **Figure 2**, gonads were fixed with 3% paraformaldehyde in 100 mM K_2_HPO_4_ for 10 minutes and permeabilized with 0.2% Triton-X. Fixed, permeabilized gonads were subsequently incubated with primary antibodies overnight at 4°C [rat anti-OLLAS (1:1000, L2 clone, Novus Biologicals, Centennial, CO, #NBP1-96713), rabbit anti-V5 (1:1000, Novus Biologicals, Centennial, CO, #NB600-381), mouse anti-FLAG (1:1000, M2 clone, Sigma, St. Louis, MO, #F3165)]. They were then incubated with secondary antibodies for 1 hour at room temperature [donkey Alexa 488 anti-rat (1:1000, Invitrogen, Carlsbad, CA, #A21208), goat Alexa 555 anti-rabbit (1:1000, Invitrogen, Carlsbad, CA, #A21429); or donkey Alexa 647 anti-mouse (1:1000, Invitrogen, Carlsbad, CA, #A31571)]. DAPI (0.5 ng/μl) was also included in the secondary antibody incubation to visualize DNA. For PGL-1∷SNAP protein imaging, gonads were incubated with 30 nM SNAP JF 549 ligand (Grimm et al. 2015) for one hour at room temperature, as previously performed (Aoki et al. 2019). Samples were mounted on glass slides with Vectashield (Vector Laboratories; Burlingame, CA)]. All staining experiments were performed a minimum of two times and yielded comparable results.

Images were captured on a Leica TCS SP8 scanning laser confocal microscope running LAS X software (version 3.5.2.18963; Leica Microsystems CMS GmbH., Buffalo Grove, IL) with a 40x oil-immersion objective and 1-1.2× zoom. Image slices (1.5 μm) were taken in sequence. Brightness and contrast were adjusted linearly and identically across all samples in FIJI/ImageJ (Schindelin et al. 2012). To quantify fluorescent signal, image stacks were z-projected by average intensity in ImageJ. Gaussian-blurred, DAPI fluorescence was used to set a threshold and mask the germlines and multiplied through the GFP and mCherry channels. Intensity of the antibody fluorescent signal was measured and averaged in 1 μm bins from the distal end of the gonad along the distal-proximal axis. For the box plot graphs, germlines were quantitated at the distal end (5-105 μm), the region where the detected reporter protein signal was strongest (**Figure S2**). Samples from at least two independent replicates were analyzed together after normalizing to a common control sample. All analysis was done using FIJI and automated using Python. Student t-tests were performed in Prism9 (version 9.0.0).

### Immunoblots

For immunoblots, young adults (18-20 hours past mid-L4 at 25°C) were boiled in 5x SDS sample buffer (250 μM Tris pH 6.8, 25 μM EDTA, 25% glycerol (V/V), 5% SDS, 70 mM 2-mercaptoethanol) for 5 minutes, analyzed by SDS-PAGE using a 10% stacking gel, transferred to a polyvinylidene difluoride (PVDF) membrane, and blocked with 5% powdered milk + PBS-T (137 mM NaCl, 2.7 mM KCl, 10 mM Na_2_HPO_4_, 1.8 mM KH_2_PO_4_, 0.1% Tween-20). Blots were incubated in 5% powdered milk + PBS-T with primary antibodies [rat anti-OLLAS (1:1000, L2 clone, Novus Biologicals, Centennial, CO, #NBP1-96713), mouse anti-V5 (1:3000, Bio-Rad, Hercules, CA #MCA1360), mouse anti-FLAG (1:1000, M2 clone, Sigma, St. Louis, MO, #F3165), rabbit anti-SNAP (1:1000, polyclonal, New England Biolabs, Ipswich, MA, P9310S), or mouse anti-tubulin (1:40,000, Sigma, St. Louis, MO, #T5168)] overnight at 4°C. After washing with PBS-T, blots were incubated with Horseradish peroxidase (HRP)-conjugated antibodies [donkey anti-mouse (H+L) (1:10,000, Jackson ImmunoResearch, West Grove, PA), goat anti-rat (H+L) (1:10,000, Jackson ImmunoResearch, West Grove, PA), or goat anti-rabbit (H+L) (1:10,000, Jackson ImmunoResearch, West Grove, PA)] in 5% powdered milk + PBS-T, washed with PBS-T, and developed with a combination of SuperSignalTM West Pico Sensitivity substrate (Thermo Scientific, Waltham, MA, #34080) and SuperSignalTM West Femto Sensitivity substrate (Thermo Scientific, Waltham, MA, #34095). Blots were imaged on Carestream Kodak BioMax light film (Sigma-Aldrich, St. Louis, MO, #1788207). The film was developed using a Konica Minolta SRX-101A film processor. Immunoblots were stripped and reprobed with western blot stripping buffer (Thermo Scientific (Pierce), Waltham, MA, #21059).

### Genetic assay for NOS-3 biological function

*nos-3* and *gld-2* single null mutants enter meiosis normally, but *gld-2(ø);nos-3(ø)* double mutants produce a synthetic germline tumor (Eckmann et al. 2004; Hansen et al. 2004). To test the function of λN-tagged NOS-3, we DAPI-stained worms of three genotypes: *(1) gld-2(q497); nos-3(q902), (2) gld-2(q497),* and *(3) gld-2(q497); nos-3(q650),* and scored by compound microscopy for presence of germline tumors (**Figure S1C**). For staining, worms were fixed with 1% paraformaldehyde in 100 mM K_2_HPO_4_ and stained with DAPI (0.5 ng/μl).

### RNA staining, imaging, and quantitation

smFISH probe sets were designed (https://www.biosearchtech.com/support/education/stellaris-rna-fish) and synthesized commercially with conjugated fluorophores by Stellaris/Biosearch Technologies (Novato, CA) (**Table S4**). The probe set directed against *gfp* exons contained 38 unique oligonucleotides labeled with CAL Fluor Red 610. The probe set against *mCherry* exons contained 39 unique olignucleotides labeled with Quasar 570. A 250 μM stock concentration of each probe set was made by dissolving lyophilized probes in RNase-free TE buffer (10 mM Tris-HCl, 1 mM EDTA, pH 8.0). A final concentration of 0.05 μM was used for hybridization of both probe sets.

Samples were prepared, stained, and imaged for smFISH as described (Lee et al. 2016). Briefly, animals were grown at 25°C for 18-20 hours past the mid-L4 stage, and their gonads extruded in PBS-T + 0.25 mM levamisole. Samples were then fixed in 3.7% formaldehyde (Fisher Scientific, Waltham, MA, F79-500) for 13-15 minutes and permeabilized in RNAse-free PBS (137 mM NaCl, 2.7 mM KCl, 10 mM Na_2_HPO_4_, 1.8 mM KH_2_PO_4_) + 0.1% Triton-X for 10-12 minutes. Samples were incubated at room temperature in PBS-T for 30-45 minutes, equilibrated in smFISH wash buffer [2x SSC (Thermo Fisher Scientific, Waltham, MA, AM9763), 10% Formamide, DEPC water, 0.1% Tween-20] for 15-20 minutes, and incubated in hybridization buffer plus *gfp* and *mCherry* smFISH exon probes (0.05 μM) at 37°C for 46-48 hours. Samples were then washed with smFISH wash buffer and DAPI (1 μg/μl) at 37°C for 40-50 minutes. Finally, samples were resuspended in 18 μl Prolong Glass mounting medium (Life Technologies Corporation, Carlsbad, CA), mounted on glass slides, and cured in the dark for at least 24 hours before imaging.

Germlines were imaged on a Leica TCS SP8 laser scanning confocal microscope equipped with a Leica HC PL APO CS2 63×/1.40 NA oil immersion objective, sensitive detectors (HyDs), standard Photomultipliers (PMTs), and LAS software (version 3.5.2.18963; Leica Microsystems CMS GmbH., Buffalo Grove, IL). Since the molecular effects of NOS-3 tethering were observed broadly in the germline, gonads were imaged in the mid-pachytene region (**Figure 4A**). The zoom factor was set to 3.0 (300% zoom), the pinhole to 95.5 μm, the window to 1024×512 pixels, and gonads were imaged at full depth with a z-step size of 0.3 μm. Images were taken using bidirectional scan at 8,000 Hz. Channels were imaged sequentially between stacks. The mCherry mRNA probed with Quasar 570 was excited with 561 nm wavelength (3.5%, HeNe) and signal collected on a HyD detector from 564-588 nm with gain set to 80. Line scans were averaged 16 times, and frames were accumulated 4 times. The GFP mRNA probed with Cal Fluor 610 was excited with 594 nm wavelength (3.5%, HeNe) and signal was collected on a HyD detector from 600-680 nm with gain set to 80. Line scans were averaged 32 times, and frames were accumulated twice. The native fluorescence of fixed GFP was excited with 488 nm wavelength (0.75%, Argon (25%)) and signal collected on a HyD detector from 495-555 nm with gain set to 80. Line scans were averaged 16 times, and frames were accumulated twice. Nuclear signal from DAPI was excited at 405 nm (1.0%, UV) and signal was collected on a PMT detector from 412-508 nm with gain set to 500-575 and an offset of −2.0%. Lines were averaged 6 times, and frames accumulated twice. Representative images were created using ImageJ. Partial maximum intensity projections were created and brightness adjusted in ImageJ. All images were treated identically.

Confocal smFISH images were analyzed by Imarisx64 (Imaris, version 9.3.1) and ImarisFileConverterx64 (Imaris, version 9.2.1) on a Dell Precision 5820 with a 64-bit Windows 10 Education operating system, an Intel(R) Xeon(R) W-1245 CPU @3.70GHz processor, and 128 GB of RAM. To count the number of signals detecting GFP and mCherry mRNAs in each image, “Spots” were identified by the software Creation Wizard for the *gfp* and *mCherry* mRNA image channels. The same framework was used for all Spots algorithms, though the filter thresholds differed between *gfp* and *mCherry* mRNA, and also from replicate set to replicate set. The general Spots Creation algorithm is as follows: Enable Region Of Interest = false; Enable Tracking = false; Source Channel Index = 1; Estimated XY Diameter = 0.250 um; Estimated Z Diameter = 0.600 um; Background Subtraction = true. The different thresholds for the [Classify Spots] “Quality” filters are as follows: *gfp* experiments 1 and 2: 4.1530; *mCherry* experiment 1: 12.922; *mCherry* experiment 2: 8.1280. The different thresholds for the [Classify Spots] “Intensity Mean” are as follows: *gfp* experiments 1 and 2: between 16.342 and 62.617; *mCherry* experiment 1: between 41.252 and 87.664; *mCherry* experiment 2: between 26.791 and 114.70. Manual thresholds were determined by eye. Identical *gfp* or *mCherry* mRNA thresholds were applied to all *gfp* or *mCherry* images within the same experimental replicate. Student t-tests were performed in Prism9 (version 9.0.0). Data from two independent experiments was combined before statistical tests were run.

---

**Table S1.** Statistics comparing fluorescent protein ratios in NOS-3 deletion mutants.

| **strain** | **v. strain** | **p-value** |
| --- | --- | --- |
| N2 wild-type | NOS-3:FLAG; dual reporter | <0.0001 |
| " | NOS-3:FLAG:λN; dual reporter | <0.0001 |
| " | NOS-3(Δ183-821):FLAG:λN; dual reporter | <0.0001 |
| " | NOS-3(Δ431-821):FLAG:λN; dual reporter | <0.0001 |
| " | NOS-3(Δ674-821):FLAG:λN; dual reporter | <0.0001 |
| " | NOS-3(Δ2-186):FLAG:λN; dual reporter | <0.0001 |
| " | NOS-3 (822 STOP):FLAG:λN; dual reporter | <0.0001 |
| NOS-3:FLAG; dual reporter | NOS-3:FLAG:λN; dual reporter | <0.0001 |
| " | NOS-3(Δ183-821):FLAG:λN; dual reporter | <0.0001 |
| " | NOS-3(Δ431-821):FLAG:λN; dual reporter | <0.0001 |
| " | NOS-3(Δ674-821):FLAG:λN; dual reporter | <0.0001 |
| " | NOS-3(Δ2-186):FLAG:λN; dual reporter | <0.0001 |
| " | NOS-3 (822 STOP):FLAG:λN; dual reporter | <0.0001 |
| NOS-3:FLAG:λN; dual reporter | NOS-3(Δ183-821):FLAG:λN; dual reporter | <0.0001 |
| " | NOS-3(Δ431-821):FLAG:λN; dual reporter | <0.0001 |
| " | NOS-3(Δ674-821):FLAG:λN; dual reporter | 0.5735 |
| " | NOS-3(Δ2-186):FLAG:λN; dual reporter | 0.4646 |
| " | NOS-3 (822 STOP):FLAG:λN; dual reporter | <0.0001 |
| NOS-3(Δ183-821):FLAG:λN; dual reporter | NOS-3(Δ431-821):FLAG:λN; dual reporter | <0.0001 |
| " | NOS-3(Δ674-821):FLAG:λN; dual reporter | <0.0001 |
| " | NOS-3(Δ2-186):FLAG:λN; dual reporter | <0.0001 |
| " | NOS-3 (822 STOP):FLAG:λN; dual reporter | <0.0001 |
| NOS-3(Δ431-821):FLAG:λN; dual reporter | NOS-3(Δ674-821):FLAG:λN; dual reporter | <0.0001 |
| " | NOS-3(Δ2-186):FLAG:λN; dual reporter | <0.0001 |
| " | NOS-3 (822 STOP):FLAG:λN; dual reporter | <0.0001 |
| NOS-3(Δ674-821):FLAG:λN; dual reporter | NOS-3(Δ2-186):FLAG:λN; dual reporter | 0.1011 |
| " | NOS-3 (822 STOP):FLAG:λN; dual reporter | <0.0001 |
| NOS-3(Δ2-186):FLAG:λN; dual reporter | NOS-3 (822 STOP):FLAG:λN; dual reporter | <0.0001 |

**Table S2:** Strains examined in this study.

| Strain | Genotype | Citation |
| --- | --- | --- |
| N2 | wild type, Bristol strain | (Brenner 1974) |
| JK3026 | *gld-2(q497)/ hT2[qIs48](I;III)* | (Eckmann et al. 2004) |
| JK3294 | *gld-2(q497)/ ccls4251 unc-15(e73) I; nos-3(q650)/ mIn1[mls14 dpy-10(e128)] II* | (Eckmann et al. 2004) |
| JK6139 | *nos-3(q940)[3xFLAG::nos-3] II; qSi380[Pmex-5::eGFP::his-58::MODC PEST::tbb-2 3'UTR 3xboxb::gpd-2::mCherry::his-58::MODC PEST::tbb-2 3'UTR mutant 3xboxb::tbb-1 intergenic] IV* | This Work |
| JK6140 | *nos-3(q902)[nos-3::λN::3xFLAG] II; qSi380[Pmex-5::eGFP::his-58::MODC PEST::tbb-2 3'UTR 3xboxb::gpd-2::mCherry::his-58::MODC PEST::tbb-2 3'UTR mutant 3xboxb::tbb-1 intergenic] IV* | This Work |
| JK6145 | *gld-2(q497)/ ccls4251 unc-15(e73) I; nos-3(q902)/ mIn1[mls14 dpy-10(e128)] II* | This Work |
| JK6268 | *qSi380[Pmex-5::eGFP::his-58::MODC PEST::tbb-2 3'UTR 3xboxb::gpd-2::mCherry::his-58::MODC PEST::tbb-2 3'UTR mutant 3xboxb::tbb-1 intergenic] IV* | This Work |
| JK6378 | *nos-3(q1176)[nos-3 (822 STOP\*)::λN::3xFLAG] II; qSi380[Pmex-5::eGFP::his-58::MODC PEST::tbb-2 3'UTR 3xboxb::gpd-2::mCherry::his-58::MODC PEST::tbb-2 3'UTR mutant 3xboxb::tbb-1 intergenic] IV. \*stop codon added after amino acid 822* | This Work |
| JK6438 | *nos-3(q1173)[nos-3 (aa Δ674-821)::λN::3xFLAG] II; qSi380[Pmex-5::eGFP::his-58::MODC PEST::tbb-2 3'UTR 3xboxb::gpd-2::mCherry::his-58::MODC PEST::tbb-2 3'UTR mutant 3xboxb::tbb-1 intergenic] IV* | This Work |
| JK6439 | *nos-3(q1174)[nos-3 (aa Δ431-821)::λN::3xFLAG] II; qSi380[Pmex-5::eGFP::his-58::MODC PEST::tbb-2 3'UTR 3xboxb::gpd-2::mCherry::his-58::MODC PEST::tbb-2 3'UTR mutant 3xboxb::tbb-1 intergenic] IV* | This Work |
| JK6440 | *nos-3(q1175)[nos-3 (aa Δ183-821)::λN::3xFLAG] II; qSi380[Pmex-5::eGFP::his-58::MODC PEST::tbb-2 3'UTR 3xboxb::gpd-2::mCherry::his-58::MODC PEST::tbb-2 3'UTR mutant 3xboxb::tbb-1 intergenic] IV* | This Work |
| JK6442 | *nos-3(q1177)[nos-3 (aa Δ2-186)::λN::3xFLAG] II; qSi380[Pmex-5::eGFP::his-58::MODC PEST::tbb-2 3'UTR 3xboxb::gpd-2::mCherry::his-58::MODC PEST::tbb-2 3'UTR mutant 3xboxb::tbb-1 intergenic] IV* | This Work |
| JK6443 | *nos-3(q902)[nos-3::λN::3xFLAG] II; pgl-1(q980)[pgl-1::3xboxb] IV* | This Work |
| JK6444 | *nos-3(q940)[nos-3::3xFLAG] II; pgl-1(q980)[pgl-1::3xboxb] IV* | This Work |

**Table S3:**
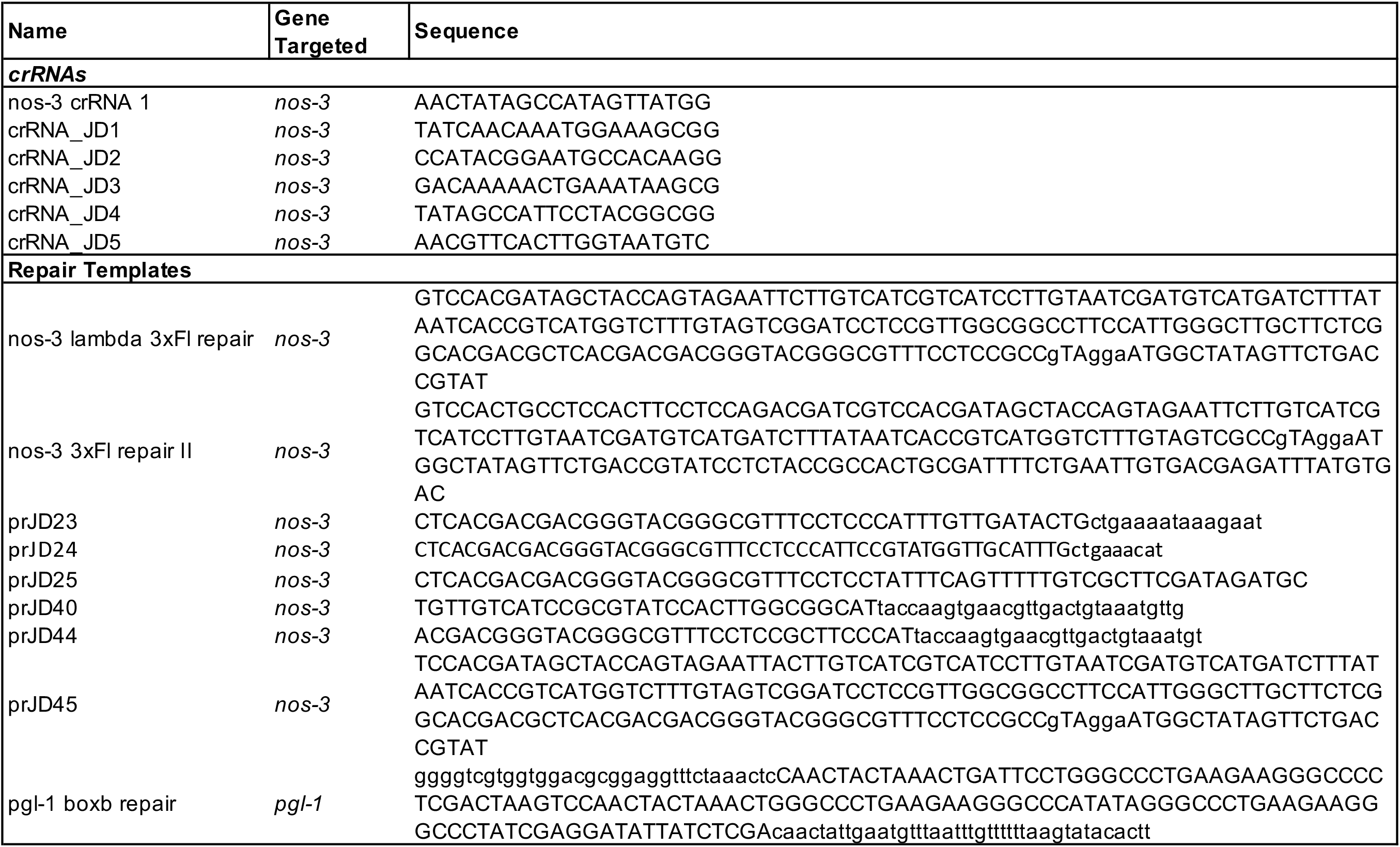
CRISPR RNAs and repair templates.

**Table S4:**
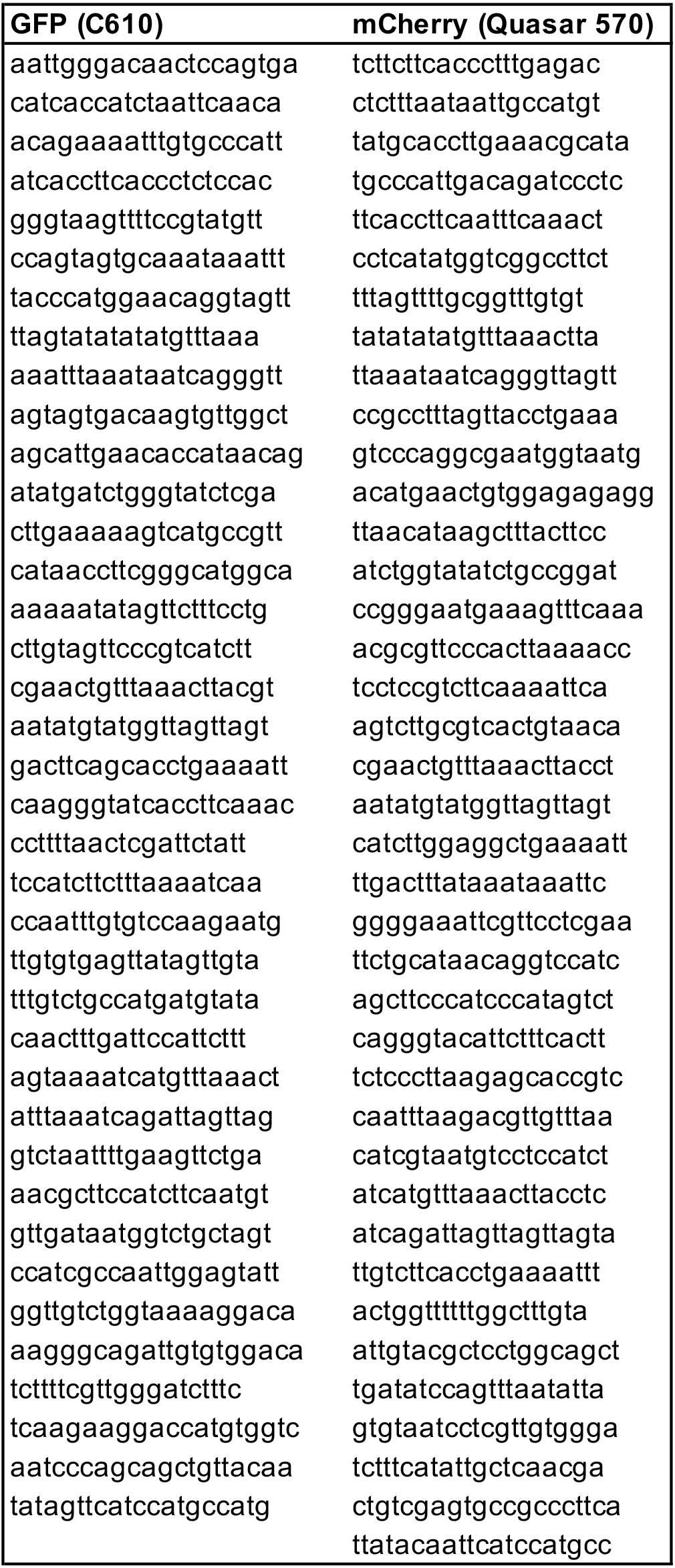
smFISH probes.

**Figure S1.**
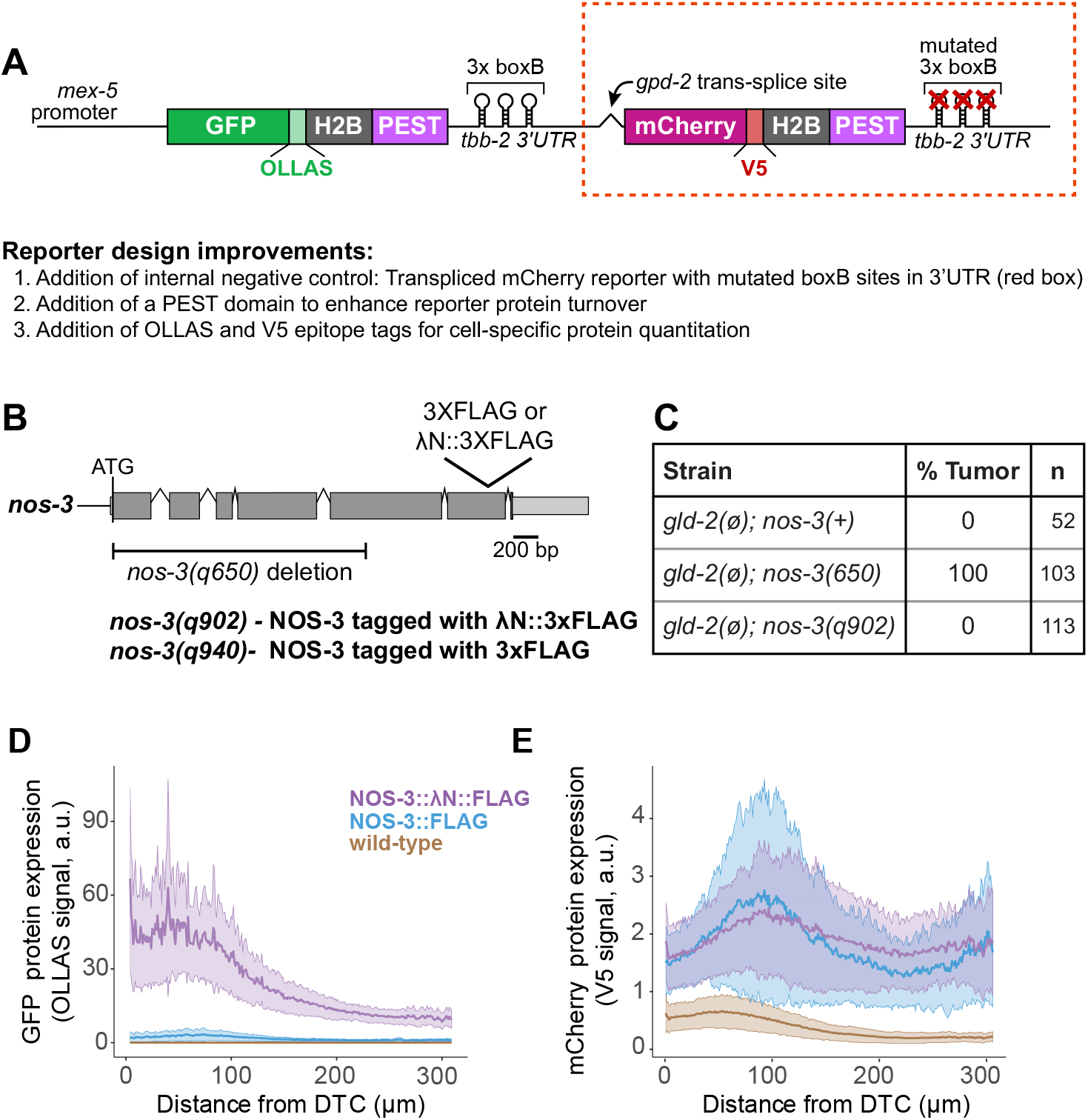
Supplemental data showing that tethered NOS-3 enhances reporter expression. **(A)** Schematic of the bicistronic GFP and mCherry dual reporter. An adult pan-germline promoter (*mex-5*) drives expression of one nascent transcript that resolves with trans-splicing to generate two mRNAs: one encoding GFP∷OLLAS∷histone H2B∷PEST domain∷3x boxB∷*tbb-2* 3’UTR and the other encoding mCherry (mCh)∷V5∷histone H2B∷PEST domain∷mutant 3x boxB∷*tbb-2* 3’UTR. The two transcripts are separated by a *gpd-2* trans-splice site. *gfp* and *mCherry* contain introns (not depicted) and that smFISH probes targeted the *gfp* or *mCherry* coding regions only. Improvements from the previous GFP tethering reporter (Aoki et al. 2018) are noted. **(B)***nos-3* locus, showing extent of the *q650* deletion and location of inserts of either FLAG or λN∷FLAG. Coding regions in dark gray; untranslated regions in light grey. **(C)** Test of tagged NOS-3 functionality in germline development. No single mutants lacking either *nos-3* or *gld-2* alone have a germline tumor (Kadyk and Kimble 1998; Kraemer et al. 1999), but a *nos-3*; *gld-2* double mutant has a germline tumor (Eckmann et al. 2004; Hansen et al. 2004). The *nos-3(q902)* allele, where NOS-3 is tagged with λN∷FLAG, behaves like wild-type *nos-3* when *gld-2* is lacking. n, number animals scored. **(D-E)** Quantitation of (D) GFP (λOLLAS) and (E) mCherry (λV5) reporter proteins, as a function of germline position in region bracketed in **Figure 1C**. Lines show averages and shading shows one standard deviation (see Methods). Wild-type serves as a negative control. Number animals scored: wild-type (n=54), NOS-3∷FLAG (n=34), NOS-3∷λN∷FLAG (n=52). The same measurements are used to generate graphs in **Figures 1E, 3D,** and **Figure S2**.

**Figure S2.**
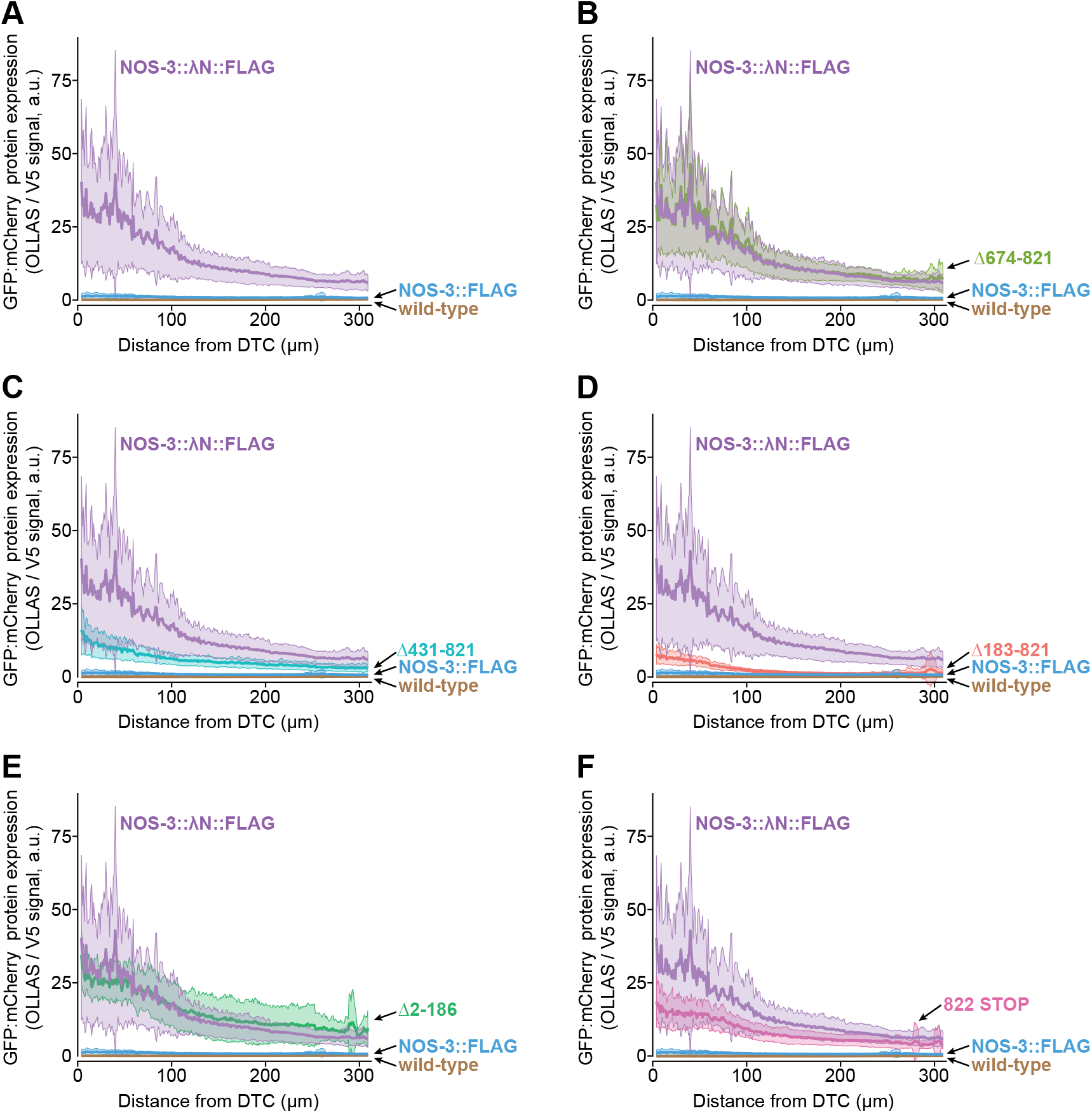
Supplemental data showing effects of tethered NOS-3 variants on reporter protein expression. Quantitation of GFP (λOLLAS) normalized to mCherry (λV5) reporter proteins, as a function of germline position in region bracketed in **Figure 1C**. Lines represent averages and shading shows one standard deviation (see Methods). Wild-type serves as a negative control. Quantitation begins at the distal end and goes for 310 μm, stopping near the proximal end of the pachytene region. N2 wild-type serves as a negative control. Each graph reports the ratio of GFP:mCherry protein expression (OLLAS:V5 signal) in **(A)** Wild-type (n=54), NOS-3∷FLAG (n=34), or NOS-3∷λN∷FLAG (n=52) expressing germlines, along with germlines expressing **(B)** NOS-3(Δ679-821)∷λN∷FLAG (n=42), **(C)** NOS-3(Δ431-821)∷λN∷FLAG (n=36), **(D)** NOS-3(Δ183-821)∷λN∷FLAG (n=37), **(E)** NOS-3(Δ2-186)∷λN∷FLAG (n=31), and **(F)** NOS-3(822 STOP)∷λN∷FLAG (n=37). Same curves and values are included in **Figures 1E, 3D** and **Figure S1D,E**.

